# Identifying repurposed drugs with moderate anti-influenza virus activity through computational prioritization of drug-target pairs

**DOI:** 10.1101/2023.07.31.551116

**Authors:** Biruhalem Taye, Roland Thünauer, Richard J Sugrue, Sebastian Maurer-Stroh, Jan Kosinski

## Abstract

Influenza A virus (IAV) causes up to five million cases of severe illness and half a million deaths worldwide each year. While there are a few clinically approved drugs for treating IAV, they are challenged by the rapid evolution of the virus leading to emergence of drug resistance and the adverse effects of the drugs. Targeting host cellular factors that support virus replication could limit resistance, increase the broad-spectrum antiviral properties of drugs, and benefit from repurposing drugs already existing against those factors. However, selecting the right drug-target pairs with low toxicity and minimal adverse effects has been challenging, even though hundreds of cellular host factors have been identified. In this study, we applied a computational and knowledge-based drug-target prioritization approach to identify promising drug-target pairs. We selected five pairs for experimental validation: telmisartan-Angiotensin II receptor, type 1 (AGTR1), metoclopramide hydrochloride-Cholinergic receptor muscarinic 1 (CHRM1), cefepime hydrochloride-phosphogluconate dehydrogenase (PGD), ranolazine dihydrochloride-sodium channel voltage-gated type v alpha subunit (SCN5A), and ofloxacin-topoisomerase II alpha 170kDa (TOP2A). Except for cefepime hydrochloride, all four drugs showed significant plaque reduction in Madin Darby canine kidney (MDCK) cells. In the immunofluorescence assay, metoclopramide hydrochloride, ranolazine dihydrochloride, and telmisartan showed antiviral activity in MDCK and/or adenocarcinoma human alveolar basal epithelial (A549) cell lines. In conclusion, our approach can prioritize and identify drugs with antiviral activity against influenza virus. Refining and strengthening such approaches could be valuable for rapid antiviral discovery and pandemic preparedness.

**Highlights:**

- Computational drug-target prioritization indicated drugs for validation
- Telmisartan showed anti-influenza virus activity in MDCK cells
- Metoclopramide and ranolazine showed anti-influenza activity in A549 and MDCK cells

## 1. Introduction

Influenza A virus (IAV) is a negative-sense RNA virus with an eight-segmented genome. It can cause pandemics, seasonal epidemics, and sporadic zoonotic infections. It is estimated that more than 50 million people died from previous pandemics, and about 0.5 million people die annually worldwide due to seasonal influenza outbreaks [1]. This number has decreased during the COVID-19 pandemic, but it is increasing again [2, 3]

IAV infects humans and a range of other animal species. The reassortment of the segmented genomes in different species can lead to antigenic shift, allowing the virus to rapidly adapt to new species and cause pandemic outbreaks [4]. Sporadic human infections with IAV strains of avian or swine origin are a major source of potential pandemics [5]. For example, throughout the summer of 2022, several outbreaks of highly pathogenic avian influenza were documented in commercial poultry farms across Europe [2]. Most of these outbreaks were caused by the H5N1 subtype, highlighting the continued threat of this virus to the poultry industry. Such outbreaks in birds and poultry worldwide raised concerns about the potential of spillover to humans [2, 6]. The emergence of new reassortant viruses with pandemic potential underscores the need for continued preparedness and response planning for emerging infectious diseases, as it has significant impact on human health and the global economy [2, 6]. Seasonal influenza viruses that circulate in humans also evolve rapidly through antigenic drift, which is caused by host immune pressure that drives it to acquire mutations to escape immune recognition [7]. The rapid evolution through the combined effects of antigenic shift and drift pose a significant challenge for influenza control [7].

Vaccination is one of the most effective strategies for mitigating influenza virus infections. However, the current seasonal influenza vaccines are not always effective and can fail to match the circulating virus strains. Additionally, the ability to develop new vaccines for upcoming pandemics on time is limited and costly. As seen with the COVID-19 pandemic, high infection rates can still occur despite vaccination. Therefore, antiviral drugs are necessary to treat infected patients and limit lockdowns and economic impact, especially at the beginning of new pandemics before vaccines are available. There are a few clinically approved antiviral drugs for influenza that target viral proteins, including M2 protein blockers (amantadine), neuraminidase inhibitors (oseltamivir, peramivir, zanamivir), and viral endonuclease polymerase acidic (PA) inhibitors (baloxavir) [8, 9]. However, the emergence of drug resistance and the adverse effects of these drugs are becoming a significant concern [8, 10]. There is, therefore, a pressing need for alternative, broad-spectrum antiviral drugs with minimal adverse effects.

The influenza virus relies on host cellular machinery for its replication. Genome-wide or targeted high-throughput siRNA or CRISPR/Cas9 screening has identified many cellular host factors (genes) that support virus replication [11–13]. Targeting these pro-viral host factors has been considered as a potential strategy for the development of anti-influenza drugs [14]. Compared to viral proteins, host factors have a minimal mutation rate, making it more difficult for the virus to develop resistance to drugs. In addition, drugs that target host factors common to different strains IAV would be more effective against new pandemic viruses. Therefore, targeting host factors can increase the broad-spectrum antiviral properties of drugs. However, targeting host factors can also increase toxicity and adverse effects by disrupting pathways essential for cell survival. Many of the identified host factors have not been characterized in follow-up studies, and it has been challenging to identify host factors that can be targeted without toxicity to cells.

Prioritization of targets among IAV cellular host factors has been limited in scope. For example, Shapira et al. (2009), Watanabe et al. (2014), and Tripathi et al. (2015) integrated siRNA screen and interactome data to prioritize targets for experimental validation [15–17]. This approach is effective at identifying promising targets, but it does not reveal how drugs against these targets would affect other cellular pathways or how to find inhibitory molecules against them. Furthermore, even if inhibitory molecules are found, most fail in clinical trials due to toxicity [18]. Drug repurposing is a strategy that has been proposed as a way to overcome issues related to toxicity and accelerate the availability of drugs with established safety profiles in the case of pandemics [18, 19]. In this approach, clinically approved drugs for other diseases that inhibit cellular host factors are evaluated for antiviral activity. Using a drug repurposing strategy, we have identified hundreds of drug-target pairs for experimental validation [11]. However, some of the clinically approved drugs, such as cancer drugs, may have side effects that are equal to or greater than influenza symptoms, indicating the need for drug prioritization. In this study, we applied computational prioritization of drugs, targets, and drug-target pairs for the selection and evaluation of drugs against the influenza virus based on their potential effectiveness and safety. The approach recapitulated drugs with known anti-influenza activity, and experimental validation *in vitro* identified four new drugs with moderate activity.

## 2. Materials and Methods

### 2.1. Drug-target prioritization

#### 2.1.1 Prioritization of drugs

Previously using network growing tools, we have identified known and potentially new host targets important for the replication of IAV based on siRNA screens datasets [11]. Then we applied knowledge-based *in silico* drug repurposing and identified 343 approved drugs [11] targeting the identified cellular host factors using MetaCore™(https://portal.genego.com/). In this study, we used those 343 drugs for prioritization. To prioritize the drugs, we used the following parameters: 1) the killing index of a drug (defined as a drug as its number of targets with serious side effects. These side effects are death, sudden death, sudden cardiac death, cardiac death, cancer, hemorrhagic strokes, heart failure and congestive heart failure) [20]; 2) the number of side effects (the predicted number of side effects of a drug are the union of side effects from all its binding targets) [20]; The side effects and killing index of each drug were obtained from the DR.PRODIS database [20]; 3) the number of influenza host factors targeted by a drug, 4) the cost of the drug, and 5) the availability on the market (number of manufacturers and packagers). The costs of the drugs and their availability were obtained from the DrugBank database [21]; 6) the likelihood ratio of a drug name co-occurrence with words of influenza virus or viruses in PubMed abstracts (see below).

#### 2.1.2 Prioritization of targets

The 349 cellular host targets of the 343 FDA approved drugs were prioritized based on three parameters: 1) the degree of a target node within the network of the cellular host targets (degree is the number of edges to other cellular targets that are connected to the target node). This was obtained by network analysis of all cellular host targets targeted by the 343 FDA approved drugs using the STRING (v10) database [22]. Highly connected targets (hubs) tend to have several cellular activities and are usually essential [23], hence, targeting such genes may impose toxicity, 2) the relation of the target with influenza or other viruses. This was obtained by calculating the likelihood of co-occurrence of the names of targets and influenza virus or viruses in the PubMed abstract (see details below). The higher the number of publications that reported target and influenza virus co-occurrence would indicate the higher confidence of the target associated with influenza virus infections. 3) The druggability of the targets. The drug-gene interaction database (DGIdb 3.0) is used to select only druggable targets [24]. The druggable targets have been defined as the genes or gene products that are known or predicted to interact with drugs, ideally with a therapeutic benefit to the patient (https://www.dgidb.org/about) [24].

#### 2.1.3 Prioritization of drug-target pairs

Text mining of the published articles in PubMed or other databases has been used to extract drug-gene relationships and automatic text mining was equal to or better than manually curated knowledge [25–27]. This text mining approach assumes that the higher the co-occurrence of two terms within the literature databases, the more significant the association is likely to be [28–30]. Hence, we hypothesise that the higher the number of publications in which the drugs and targets co-occur, the more likely the drugs are active on the targets. To quantify the drug-target associations in literature, we used online PubMed “esearch” tool and counted the number of publications related to the drugs and their corresponding targets.

To account for unequal frequency distribution of the words of the drug, the target, influenza virus or viruses in the pubmed abstracts, we estimated the co-occurrence scores as the likelihood ratio:

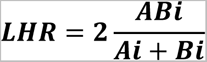

Where, LHR = Likelihood ratio, ABi = the number of PubMed abstracts that both Ai and Bi co-occur; Ai = the number of publications in Ai (e.g. drug); the number of publications in Bi (e.g. target). Separate LHR values were calculated for co-occurrences of the word drug-target, drug-influenza virus, drug-virus, target-influenza virus and target-virus associations.

#### 2.1.4. Ranking of drug-target pairs

The values of each parameter described above for drugs, targets, and drug-target pairs were standardised by calculating Z-scores:

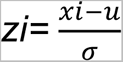

Where, Zi = Z-score, ***χ***i = value, ***μ*** = mean, ***σ*** = standard deviation.

The total Z-score was calculated by summing the values of text-mining LHRs, number of targets targeted by the drugs, number manufacturers and packagers text-mining and subtracting the Z-score values of the killing index, side effects, degree of the targets in the interaction network, and cost of the drugs for prioritising the drugs. The drug-target pairs were then ranked according the total Z-score, with the higher absolute values of the total Z-score corresponding to the more likely pairs.

### 2.2. Cells and Viruses

A549 and Vero 76 cells were maintained in DMEM media with 10% FBS, 1% Penicillin-Streptomycin, and 1% Glutamine. MDCK cells were maintained in MEM media with 10% FBS, 1% Penicillin-Streptomycin, and 1% Glutamine. The laboratory strain Influenza virus A/WSN/1933(H1N1) was generously provided by Prof. Dr Gülsah Gabriel’s at Leibniz Institute of Virology in Hamburg. The drugs telmisartan (Catalog No.: T8949, ≥ 98% (HPLC)), metoclopramide hydrochloride (Catalog No.: M0763, <= 100%), ofloxacin (Catalog No.: O8757, ≥ 99% (HPLC)), cefepime hydrochloride (Catalog No.: 1097636), ranolazine dihydrochloride (Catalog No.: R6152, ≥ 98% (HPLC)) were purchased from Sigma-Aldrich. The drugs were dissolved in Dimethyl sulfoxide (Catalog No.: D2650, ≥ 99.7%, Sigma-Aldrich) except for ofloxacin, which was dissolved in 1 mM NaOH.

### 2.3. Propagation of H1N1 virus in Vero cells

The A/WSN/1933(H1N1) virus was propagated in Vero 76 cells as described before [31]. Confluent monolayer Vero 76 cells were infected with A/WSN/1933(H1N1) at a multiplicity of infection (MOI) of 0.01 for 1 hour and the virus inoculum was washed with 1xPBS. The infected cells were incubated in 0.2% bovine serum albumin (BSA) DMEM media with 0.4μg/ml TPCK-trypsin (Catalog No.: T1426) at 37 °C in humidified 5% CO_2_ for 72 hours. The 0.4μg/ml TPCK-trypsin was added every 24 hours. The supernatant of the infected cells was centrifuged at 600 rpm for 5 minutes at 4 °C. The cell-free virus was harvested, aliquoted and stored at −80 °C. Plaque assay was used to measure the titer of the propagated virus and was 5.46×10^6^ pfu/ml. For each infection experiment, new aliquots of the virus were used.

### 2.4. Plaque and plaque reduction assay

Plaque assays were performed as described before with minor modifications [32]. Briefly, serially diluted virus was adsorbed on confluent monolayer MDCK cells in 6 well plates. After one hour of adsorption, cells were incubated with overlay media for 72 hours at 37 °C in humidified 5% CO_2_. The overlay media is a mix of an equal volume of 0.25% Avicel (RC-581 NF) in distilled water and 0.4% BSA MEM (Temin’s modification, 2X, Catalog No.: 21935028). Plaques were counted visually and the virus titer was determined as described before [33]. For the plaque reduction assay, first, the MOI was adjusted to produce 15 to 45 plaques in 6 well plates as described before [19]. Confluent monolayer MDCK cells were infected for 1 hour. The infected cells were then incubated with overlay media containing different concentrations of the drugs at 37 °C in humidified 5% CO_2_. After 72 hours of incubation, plaques were visually counted.

### 2.5. Cytotoxicity assay

The cell toxicity of the drugs was determined using Cell Counting Kit-8 (CCK-8) (Catalog No.: 96992-100TESTS-F, Sigma-Aldrich) according to the manufacturer’s directions. Briefly, 5000 MDCK cells were seeded in 96 well plates and incubated for 24 hours at 37 °C in humidified 5% CO_2_. The cells were then incubated with different concentrations of the drugs in 0.2% BSA DMEM at 37 °C in humidified 5% CO_2_. After 72 hours, cells were treated with CCK-8 solution for 2 hours. Absorbance at 450nm was measured using a microplate reader. The percentage of cell viability was calculated relative to the wells with solvent controls. The cell cytotoxicity of the drugs on A549 and MDCK cells was also determined using an immunofluorescence assay (see below). The cells were seeded at a density of 10,000 and incubated for 24 hours as described above. Then cells were treated with different concentrations of the drugs for 24 hours. Dead cells were washed off and cells attached to the wells were fixed with 3% PFA. After permeabilizing the cells with 0.1% TritonX100, cellular nuclei were stained with DAPI (4′,6-diamidino-2-phenylindole). Cell nuclei were counted as described below (Microscopy and image analysis). The percentage of cell viability was determined relative to nuclear count in solvent control wells.

### 2.6. Immunofluorescence assay

In the post-infection treatment (first scenario) either A549 or MDCK cells were infected with the virus for 1h and then cells were treated with different concentrations of the drugs. In the pre-and post-infection treatment (second scenario) A549 cells were pre-treated with different concentrations of the drugs for 8hrs and were infected with the virus for an hour. Then cells were treated with similar corresponding concentrations of these drugs for 24hrs. Briefly, about 10,000 A549 and MDCK cells were seeded in 96 well plates (CellCarrier-96 Ultra Microplates Catalog No.: 6055300, PerkinElmer) and were grown at 37 °C in humidified 5% CO_2_ for 24hrs. After washing with 1xPBS, the cells were adsorbed with interferon-free A/WSN/1933(H1N1) virus at MOI of 0.1 and were incubated for 24 hours. After removing the supernatant and washing the cells with 1xPBS, the cells were fixed with 3% paraformaldehyde for 20 minutes. The cells were permeabilized with 0.1% TritonX100 for 15 minutes and blocked with 3% BSA for 30 minutes for non-specific binding. These were followed by subsequent staining with 50 μl of primary antibody (Anti-Influenza A Virus Nucleoprotein antibody [C43] (ab128193, Abcam), 1:500 dilution) for 1 hour and 50 μl of secondary antibody (Goat anti-Mouse IgG (H+L) Highly Cross-Adsorbed Secondary Antibody, Alexa Fluor Plus 488 (Catalog No.: A32723, Thermo Fisher Scientific, 1:1000 dilution) for 1 hour in a dark humidified chamber. Fifty microliters of 1:1000 diluted 300μM DAPI (Catalog No.: D3571, Sigma-Aldrich) were used for counterstaining.

### 2.7. Microscopy and image analysis

The immunofluorescently stained cells on the 96 well plates were imaged using a Leica DMi8 inverted wide-field microscope at the Centre for Structural Systems Biology (CSSB), Advanced Light and Fluorescence Microscopy (ALFM) facility. Before imaging, two or more focal points were defined for each well. After adjusting optimum exposure times for DAPI (blue) and Alexa Fluor 488 (green) channels, the microscope was automated to take images covering each well using a 10x NA 0.32 objective. Three independent replicates of the immunofluorescence experiments and imaging were carried out for each drug. A customized Java script was used to count the number of cells (DAPI, blue) and virus-infected cells (NP, Alexa Fluor 488, green) in each image. The antiviral activity of the drugs in different concentrations was then evaluated by the percentage of virus-infected cells relative to the corresponding solvent controls. Dose-response curves fitted using Graphpad Prism v8 were used to calculate the CC_50_ and EC_50_ of the drugs.

## 3. Results

### 3.1. Prioritization of drug-target pairs

We used the systems biology and systems pharmacology approach to prioritize drugs, targets, and drug-target pairs combinations of the 343 FDA approved drugs that we previously predicted to inhibit host factors important for influenza virus replication [11]. Primarily, a total of 11 parameters were used to prioritize the drug-target pairs, such as the drug-killing index and the side effects of the drugs, cost, availability, the number of host factors targeted by the drug, the connectivity of the targets, and LHR of drug-target co-occurrence in PubMed abstracts (see Material and Methods).

The ranked drug-target pairs with calculated individual and total z-scores are described in Supplementary Data 1A. Among the 534 prioritized drug-target pairs, 399 targets were druggable (Supplementary Data 1B). The anti-hypertensive (Losartan) [34], the anti-malaria (Chloroquine) [35, 36], and the anti-influenza (Zanamivir) [37], which are known to have anti-influenza activity, are among the top-ranked drugs. High-ranking drugs with known anti-influenza activity validate the strength of our approach. For the experimental validation, we selected five high-ranking drug-target pairs with host factors involved in different stages of the IAV infection cycle and drugs that have not been tested for anti-influenza activity: telmisartan-Angiotensin II receptor, type 1 (AGTR1), metoclopramide hydrochloride-Cholinergic receptor muscarinic 1 (CHRM1), cefepime hydrochloride-phosphogluconate dehydrogenase (PGD), ranolazine dihydrochloride-sodium channel voltage-gated type v alpha subunit (SCN5A), and ofloxacin-topoisomerase II alpha 170kDa (TOP2A) (Supplementary Data 1C).

### 3.2. Anti-influenza virus activity of the selected drugs using plaque reduction assay (PRA)

First, we evaluated the cell cytotoxicity of the drugs on MDCK cell lines at 72h post-treatment (Table 1). Telmisartan and ofloxacin showed the lowest (96.01µg/ml) and highest (655.2µg/ml) CC_50_ in MDCK cells, respectively. To avoid the confounding effect of interferon, interferon-free A/WSN/1933(H1N1) virus produced in Vero 76 cells (which lack genes that encode for type I interferon [38]) was used for all experiments.

**Table 1.**
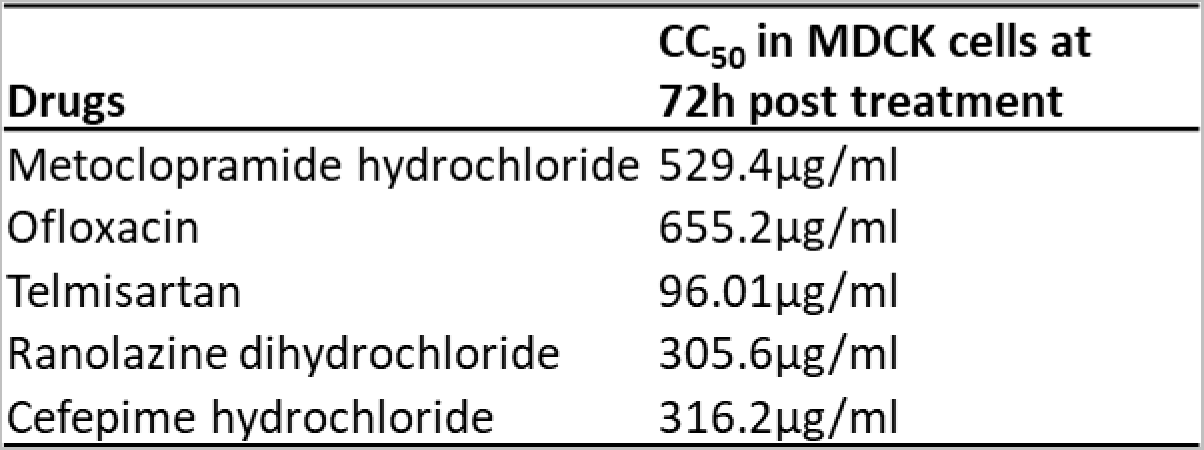
The cell cytotoxicity of the tested drugs in MDCK cells.

| **Drugs** | **CC<sub>50</sub> in MDCK cells at 72h post treatment** |
| --- | --- |
| Metoclopramide hydrochloride | 529.4µg/ml |
| Ofloxacin | 655.2µg/ml |
| Telmisartan | 96.01µg/ml |
| Ranolazine dihydrochloride | 305.6µg/ml |
| Cefepime hydrochloride | 316.2µg/ml |

The anti-influenza virus activity of the selected drugs was then measured using PRA in MDCK cells. In this assay, the plaque reduction at the non-toxic concentration of the drugs was measured relative to the solvent controls. The result showed that four of the five selected drugs significantly reduced plaque formation at the tested concentration (Fig. 1A, Supplementary Fig. 1). Following these, we determined the half-maximal effective concentration (EC_50_) and selectivity index (SI) of the four drugs that showed anti-influenza activity using PRA. As indicated in Fig. 1B, telmisartan has the lowest EC_50_ (36.6μg/ml) followed by ofloxacin (236.7μg/ml), metoclopramide hydrochloride (314.3 μg/ml), and ranolazine dihydrochloride (316μg/ml). The SI of telmisartan (2.62) is comparable with ofloxacin (2.76).

**Fig. 1.**
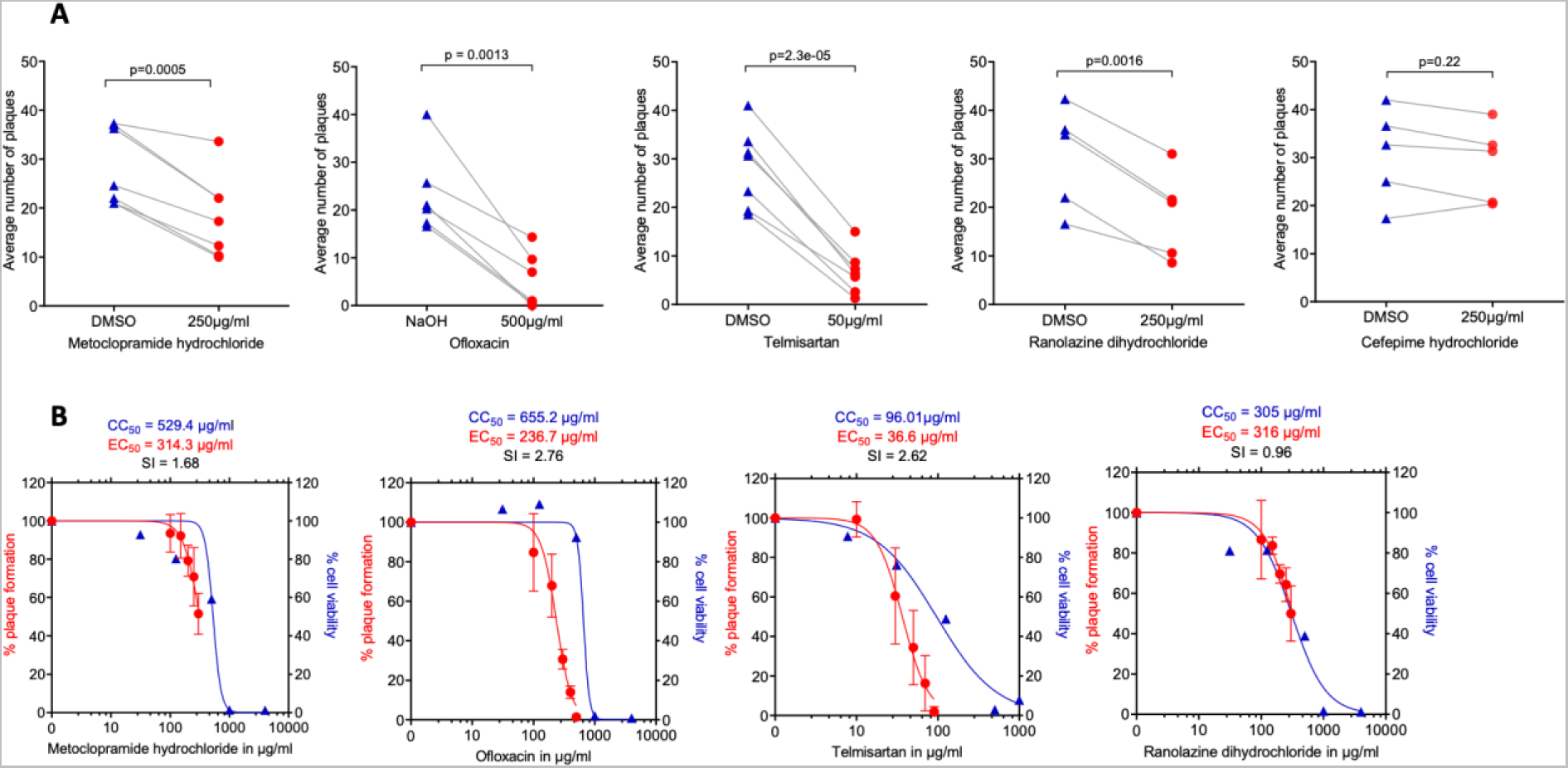
Anti-influenza virus activity of the selected drugs in MDCK cells using plaque reduction assay (PRA). A. Anti-influenza virus activity of the drugs in multiple (5-7) independent PRA experiments each with three technical replicates at a single non-toxic concentration. Each data point represents the average count of plaques from the three technical replicates. Paired t-test was used to measure the statistical significance of the differences. B. The activity of the drugs at different concentrations using three independent experiments each with three technical replicates. Dose-response curve fitting, EC_50_ and CC_50_ were performed using GraphPad Prism V8.

### 3.3. Anti-influenza activity of the selected drugs using immunofluorescence staining

Next, we evaluated the selected drugs using immunofluorescence staining, which, in contrast to PRA, allows measuring inhibition of both primary and secondary infection using human lung epithelial cells (A549). Drug or solvent control-treated virus infected cells in 96-well plates were stained with primary (mouse anti-influenza NP) and secondary (goat anti-mouse conjugated with Alexa Fluor 488) antibodies followed by counterstaining of cell nuclei with DAPI. A total of 259,200 images were obtained using high-throughput imaging from experiments of two scenarios (see Materials and Methods 2.6): 1) post-infection treatment of the drugs in A549 and MDCK cells, and 2) pre- and post-infection treatment of the drugs in A549 cells.

In the first scenario, a total of 103,680 images were analyzed. Metoclopramide hydrochloride (EC_50_ = 85.48 μg/ml) and ranolazine dihydrochloride (EC_50_ = 208 μg/ml) showed anti-influenza activity in A549 cells (Fig. 2A). Telmisartan (EC_50_ = 34.29 μg/ml), metoclopramide hydrochloride (EC_50_ = 224.7 μg/ml), and ranolazine dihydrochloride (EC_50_ = 254.9 μg/ml) showed antiviral activities in MDCK cells (Fig. 2B). In contrast, metoclopramide hydrochloride and ranolazine dihydrochloride showed antiviral activity regardless of the cell type–in both A549 and MDCK cells–with higher SI in A549 cells. Examples of the immunofluorescence images are shown in Supplementary Fig. 2.

**Fig. 2.**
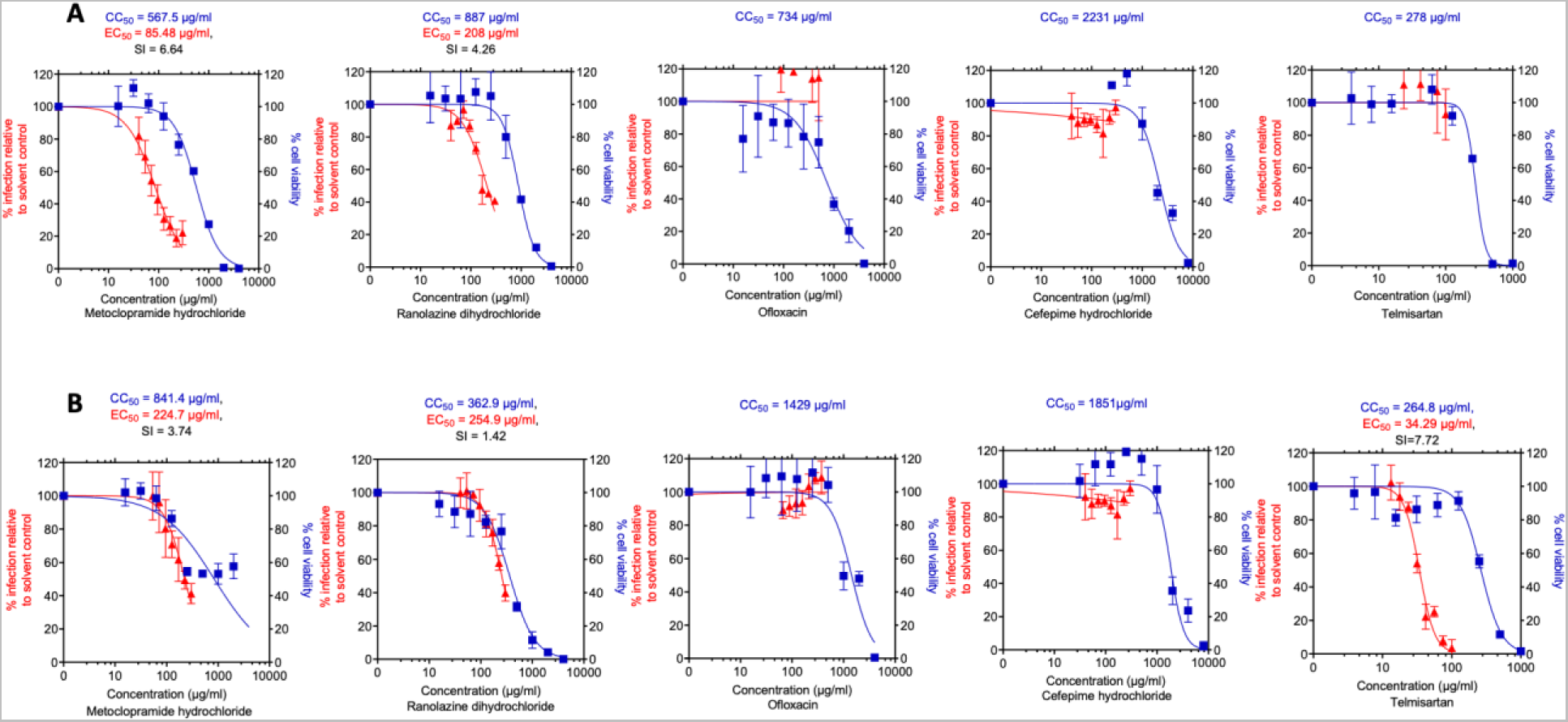
Anti-influenza A virus activity of the selected drugs in A549 and MDCK cells using immunofluorescence staining (post-infection treatment). A549 and MDCK cells were infected with A/WSN/1933(H1N1) at MOI of 0.1 for 1 hour and treated with the drugs for 24 hours. Cells were stained with anti-influenza NP and DAPI. The anti-influenza activity of the drugs in different concentrations was evaluated as the percentage of cells exhibiting NP staining relative to the solvent controls. The experiment was performed in triplicate and GraphPad Prism V8 was used for dose-response curve fitting. Each data point represents the average % of infection relative to the solvent controls. A. Anti-influenza activity of the drugs in A549 cells at increasing concentrations. B. Anti-influenza activity of the drugs in MDCK cells at increasing concentrations.

In the second scenario, A549 cells were pre-treated with different concentrations of drugs for 8 hours; after which the drugs were removed and the cells were infected with the virus for 1 hour, followed by treatment of the drugs for 24 hours with the corresponding concentrations. A total of 155,520 images were obtained from three independent experiments, each with triplicate (9 plates, 60 wells per plate, and 288 images per well). Metoclopramide hydrochloride and ranolazine dihydrochloride showed anti-influenza activity. The pre-and post-infection treatment of ranolazine dihydrochloride showed increased anti-influenza activity (EC50 = 157.1µg/ml) compared to only post-infection treatment in A549 cells (EC50 = 208 µg/ml) (Fig. 2A, Fig. 3).

**Fig. 3.**
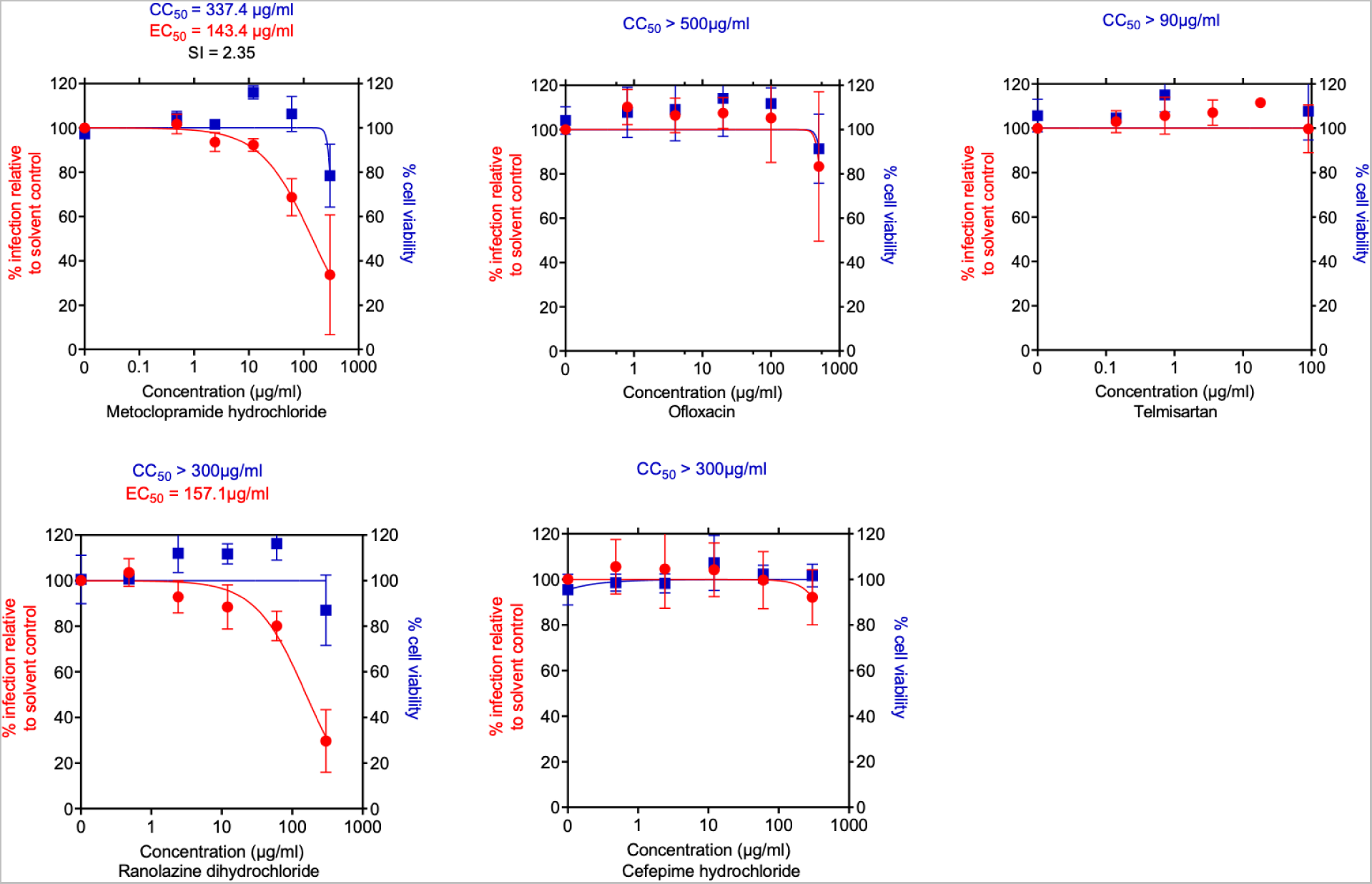
Anti-influenza virus activity of the selected drugs in A549 cells using immunofluorescence staining (pre- and post-infection treatment). A549 cells were pre-incubated with the drugs for 8 hours, infected with A/WSN/1933(H1N1) at MOI of 0.1 for 1 hour, and again treated with the drug for 24 hours. Cells were stained with anti-influenza NP and DAPI. The anti-influenza activity of the drugs in different concentrations was evaluated as the percentage of cells exhibiting NP staining relative to the solvent controls. GraphPad Prism V8 was used for dose-response curve fitting. Each data points in the graph represent the average percentage of infection relative to the solvent controls from three independent experiments, each with three technical replicates.

## 4. Discussion

The rapid evolution of influenza viruses imposes a huge challenge for the control of influenza virus either using vaccination or the few existing anti-influenza drugs. Hence, there is an urgent need for new, safe, affordable, and broad-spectrum anti-influenza drugs that resist the rapid evolution of the virus. Targeting cellular host factors that are important for virus replication could be considered one of the strategies for discovering broad-spectrum anti-influenza drugs. To this effect, there have been a dozen high-throughput screening and computational studies that identified important cellular host factors for the virus [8–13]. However, as these cellular host factors could also be involved in pathways important for the survival of the cell, one should be very cautious while targeting these targets. One of the strategies used to reduce such effects is drug repurposing. However, the side effect of newly proposed drugs against the influenza virus should not exceed influenza symptoms.

In this study, we applied systems biology and pharmacology approach for quantitative prioritization of cellular host targets, drugs, and drug-target pairs to select safe, affordable, and broad-spectrum anti-influenza drugs. This approach considered parameters on drug safety and cost, the druggability of the host factor targets, and the strength of the association between the drug and host targets in the scientific literature. We have demonstrated that this approach prioritized known and novel drugs with anti-influenza virus activity at the top of the ranking list. For instance, among the top 15 ranked drug-target pairs, Losartan [34], Chloroquine [35, 36], and Zanamivir [37] are drugs with known anti-influenza activity. Based on the knowledge of the influenza virus infection cycle and novelty, we selected five drug-target pairs for validation. Four of the five experimental validated drugs showed anti-influenza virus activity in cell culture. Telmisartan, the 4^th^ ranked drug, showed strong anti-viral activity in both plaque reduction and immunofluorescence assays in MDCK cells, while the 12^th^ ranked ranolazine and the 33^rd^ ranked metoclopramide showed anti-viral activity both in A549 and MDCK cells. Among experimentally tested drugs, ofloxacin and cefepime are ranked lowest. Indeed, cefepime has no antiviral activity, and the antiviral activity of ofloxacin is limited to PRA. Taken together, the data demonstrated a proof of concept that our prioritization approach is capable of selecting known as well as novel drugs with anti-influenza activity.

In our experimental validation, the anti-viral activities of the tested drugs were affected by the cell type and experimental assay. For instance, telmisartan is anti-viral in MDCK cells but not in A549 cells, whereas ranolazine dihydrochloride and metoclopramide hydrochloride are active in both cell lines. According to our previously published expression data [39], these differences are not due to a different expression of the corresponding targets in these cell lines, as the expression of the targets in both cell lines is comparable (Supplementary Fig. 3), prompting further investigations.

Corroborating our results, Telmisartan, metoclopramide, and ranolazine have been increasingly being studied for their antiviral properties. Telmisartan is a selective blocker of AGTR1 [41] and has been shown to have multiple cardiovascular benefits, including reducing blood pressure, inflammation, and oxidative stress through the activation of peroxisome proliferator-activated receptor gamma (PPARγ) [42]. Recent studies have demonstrated the effect of telmisartan against Chikungunya Virus (CHIKV) [44], herpes simplex virus 1 (HSV-1) [44], and Japanese encephalitis virus (JEV) [44]. Its antiviral effects against CHIKV are attributed to the direct inhibition of CHIKV nonstructural protein P2 (nsP2) protease activity and the reduction of inflammation [44, 45]. It also reduced mortality in COVID-19 patients [43, 46] and protected mice from lethal encephalitis caused by neuroadapted Sindbis virus (NSV) and western equine encephalitis virus (WEEV) [47]. Another AGTR1 blocker losartan has also been shown to reduce viral titers in the lungs and attenuate H7N9-induced lung injury in mice, suggesting both antiviral and anti-inflammatory benefits of AGTR1 blockers [34]. Further studies are needed to elucidate the antiviral mechanism of telmisartan and losartan. In addition, a retrospective or prospective study on patients who regularly take AGTR1 blockers could shed more light on their protective effect against influenza viruses.

The antiemetic metoclopramide inhibits CHRM1 [48, 49]. CHRM1 is known to strongly activate phosphatidylinositol (PI) hydrolysis and substantially elevate cyclic adenosine monophosphate (cAMP) levels [50]. The intermediate products of PI hydrolysis (PI(4,5)P_2_), elevated cAMP, and phosphoinositide 3-kinase (PI3K) pathway are important for influenza virus viral genome packaging and propagation [51–53]. Hence, we speculate that metoclopramide inhibits IAV through the reduction of PI hydrolysis. Metoclopramide was reported to have antiviral activity against Dengue virus (DENV) in vivo and in vitro mouse models, possibly through inhibition of the Dopamine 2 receptor (D2R) [54].

The increased cellular concentration of K^+^ and the switch from Na^+^ to K^+^ trigger the uncoating of influenza and other RNA-enveloped viruses through endosomal acidification [55, 56]. Thus, ranolazine, as the inhibitor of Na+ channel protein SCN5A and other channel proteins, may inhibit endosomal maturation and IAV infectivity.

This study is not without limitations. Although the tested drugs have their protein targets annotated in databases, we have not confirmed whether these targets are indeed responsible for the effects on the IAV infection. Also, as the experimentally tested drugs targeted host factors important for different stages of the influenza infection cycle, testing drug combinations could show higher synergistic antiviral activity. Finally, and most importantly, the tested drugs showed anti-viral activity only at higher concentrations, and hence these drugs are less likely to be considered as lead compounds for drug development. Still, telmisartan, ranolazine dihydrochloride, and metoclopramide hydrochloride showed anti-viral activity. Thus, might be worth investigating the prevalence and severity of influenza virus infection in patients regularly taking these drugs in their medications.

In conclusion, our quantitative drug-target pair prioritization can select known and novel drugs with detectable anti-influenza virus activity. Four out of five tested drugs showed activity, demonstrating that active drugs can be identified with a high success rate using even relatively simple bioinformatics prioritization strategies. Extending the computational pipeline with additional multi-omics and drug-target data layers might further increase the success rate and potentially identify drug-target pairs with higher activities, benefitting the rapid selection of anti-viral drugs for upcoming pandemic preparedness. Such an approach could also be extended to other rapidly evolving viral diseases.

## Acknowledgements

We thank Andrea Mikulasova for her technical support.

## Declaration of generative AI and AI-assisted technologies in the writing process

During the preparation of this work the authors used Grammarly and ChatGPT in order to improve grammar and style in the abstract, introduction, and discussion. After using this tool/service, the author(s) reviewed and edited the content as needed and take(s) full responsibility for the content of the publication.

