## Supplementry Figures for "Identifying repurposed drugs with moderate anti-influenza virus activity through computational prioritization of drug-target pairs"

### Supplementary Figures

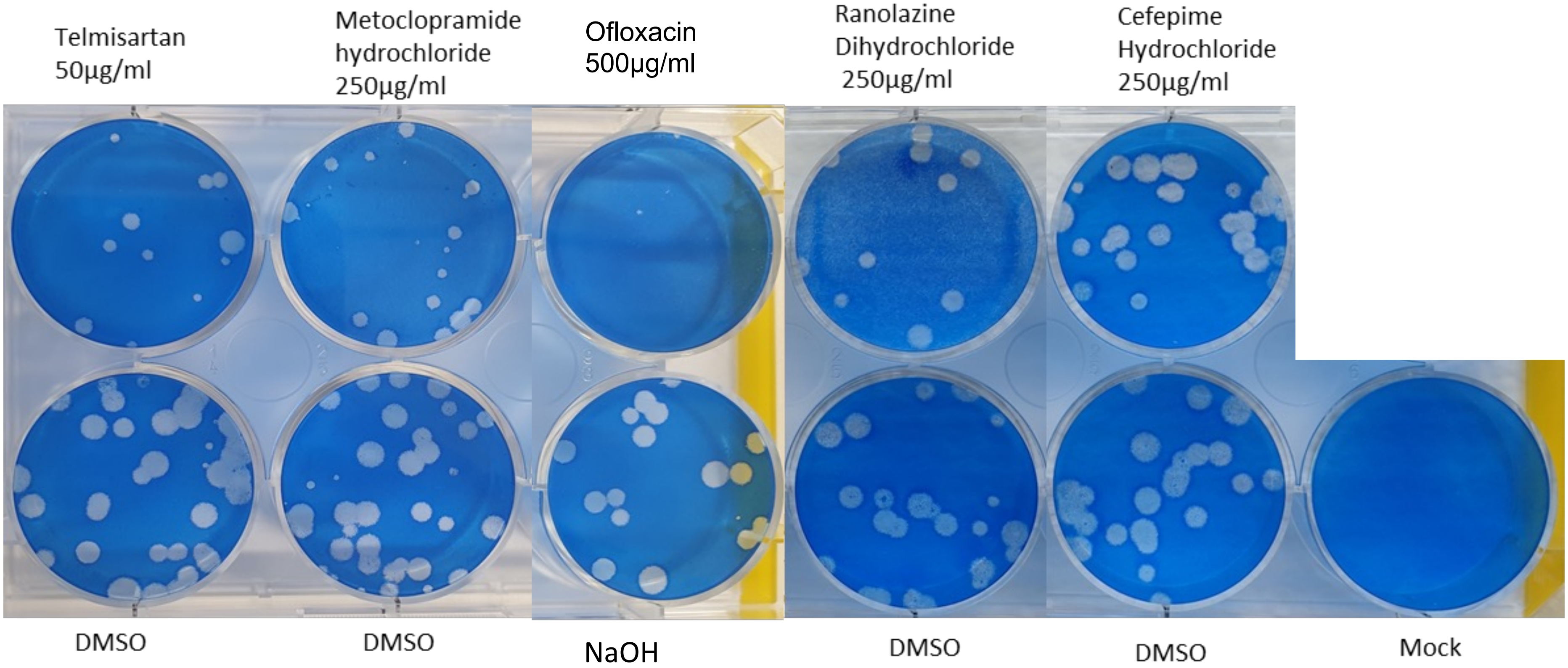


**Sup.Fig 1.** Representative plaque reduction assay (PRA) images showing the anti-influenza activity of the drugs at the tested concentrations.

A

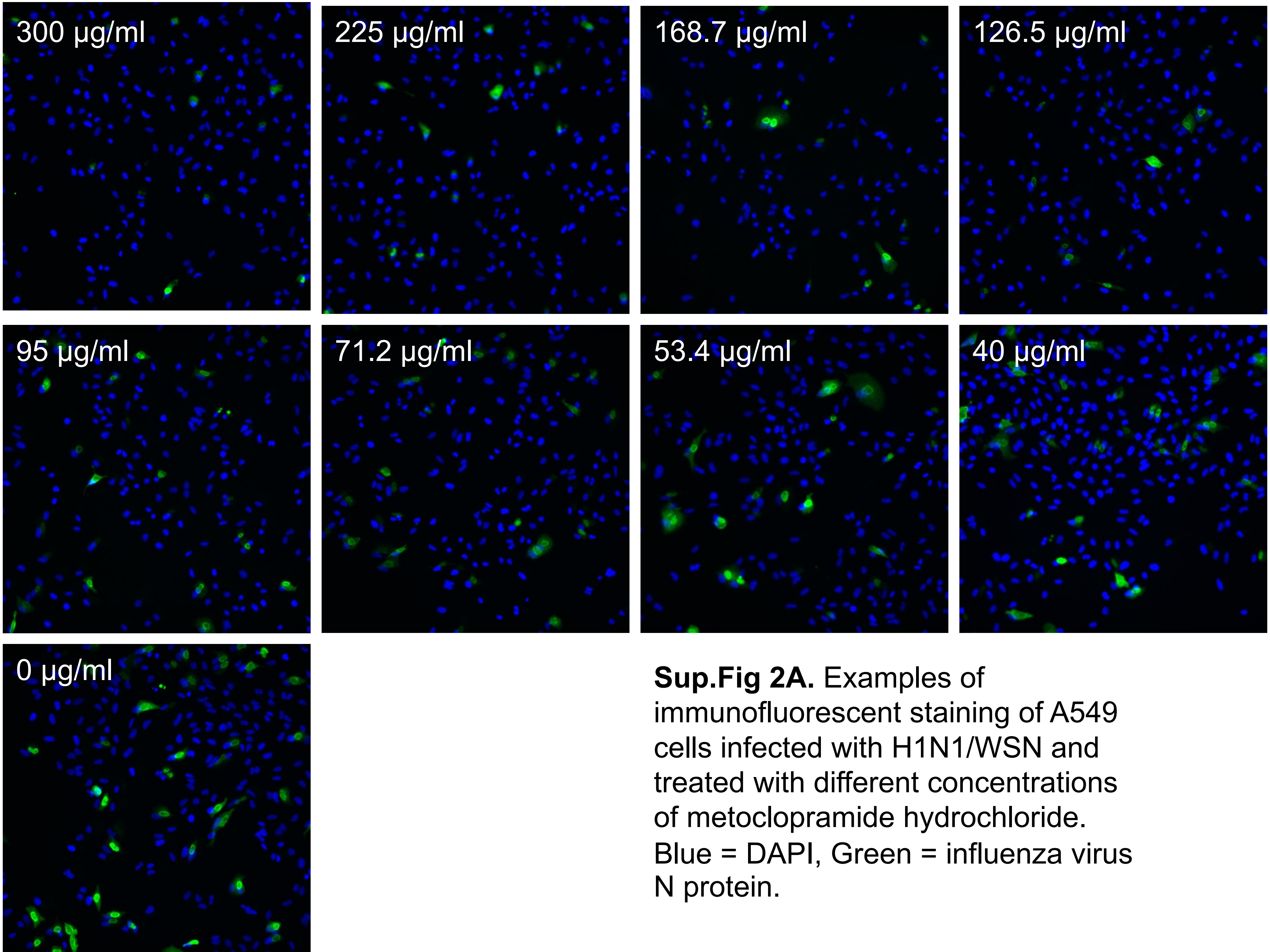


**Sup.Fig 2A.** Examples of immunofluorescent staining of A549 cells infected with H1N1/WSN and treated with different concentrations of metoclopramide hydrochloride. Blue = DAPI, Green = influenza virus N protein.

B

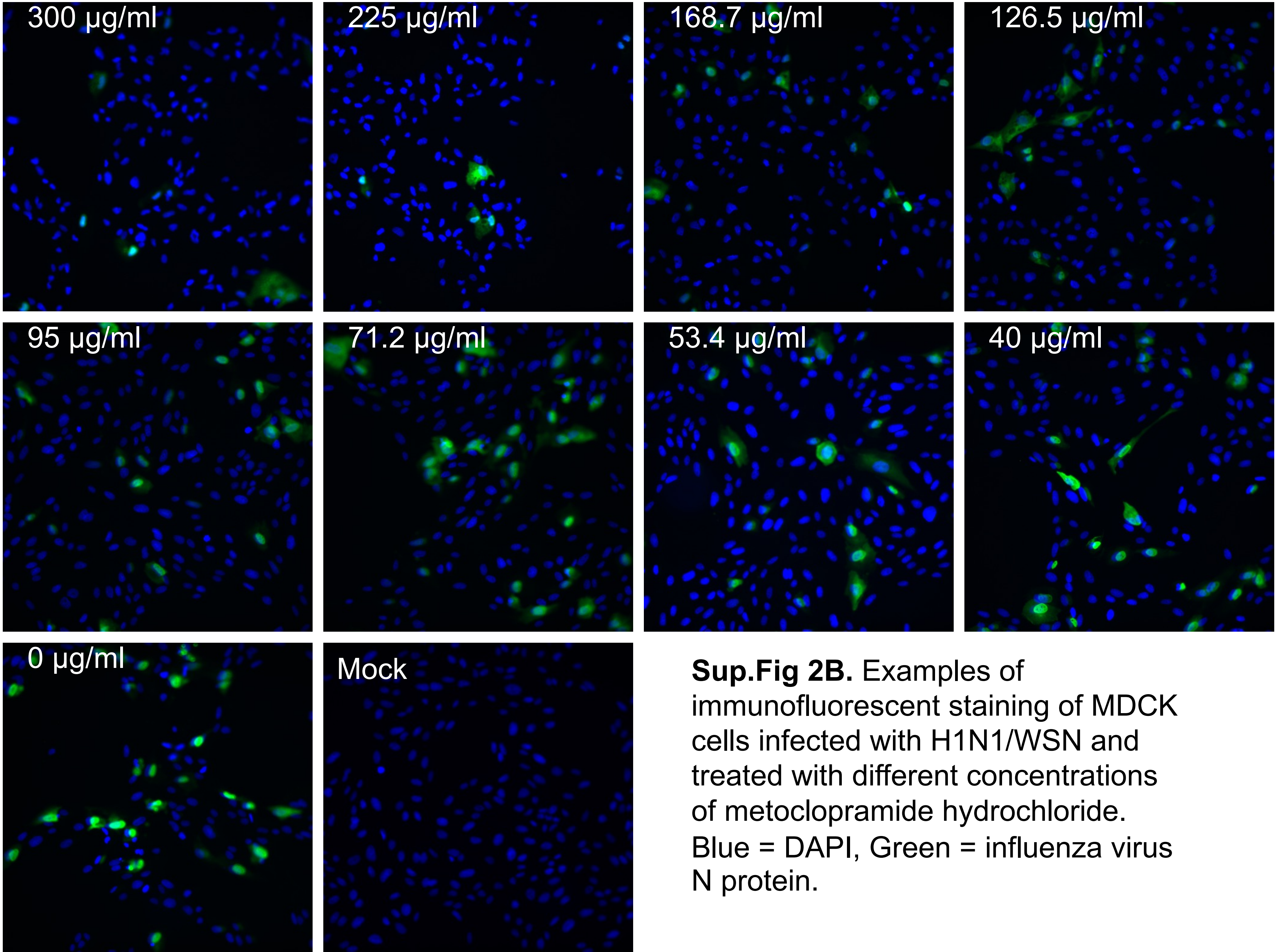


**Sup.Fig 2B.** Examples of immunofluorescent staining of MDCK cells infected with H1N1/WSN and treated with different concentrations of metoclopramide hydrochloride. Blue = DAPI, Green = influenza virus N protein.

C

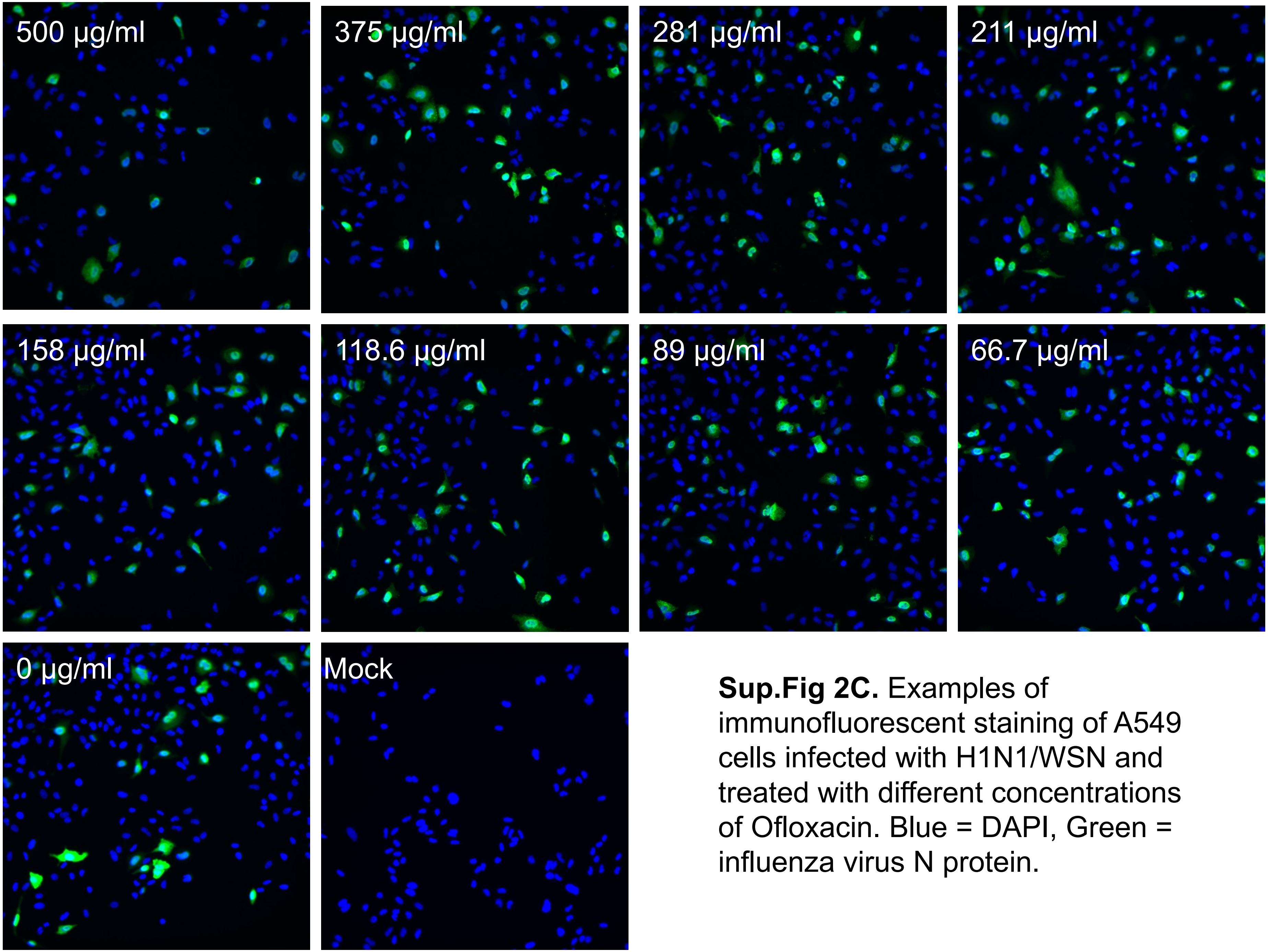


**Sup.Fig 2C.** Examples of immunofluorescent staining of A549 cells infected with H1N1/WSN and treated with different concentrations of Ofloxacin. Blue = DAPI, Green = influenza virus N protein.

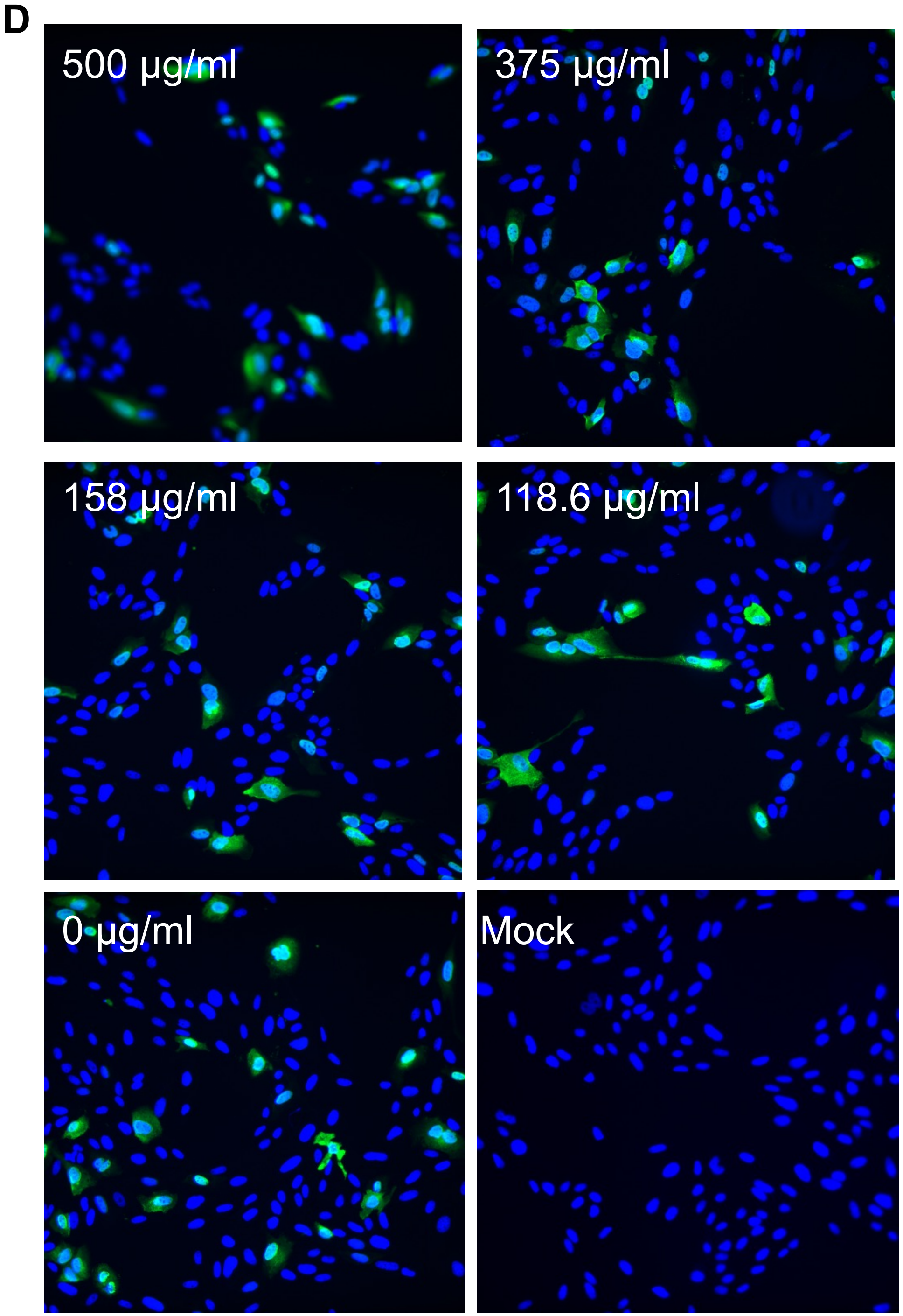


**Sup.Fig 2D.** Examples of immunofluorescent staining of MDCK cells infected with H1N1/WSN and treated with different concentrations of Ofloxacin. Blue = DAPI, Green = influenza virus N protein.

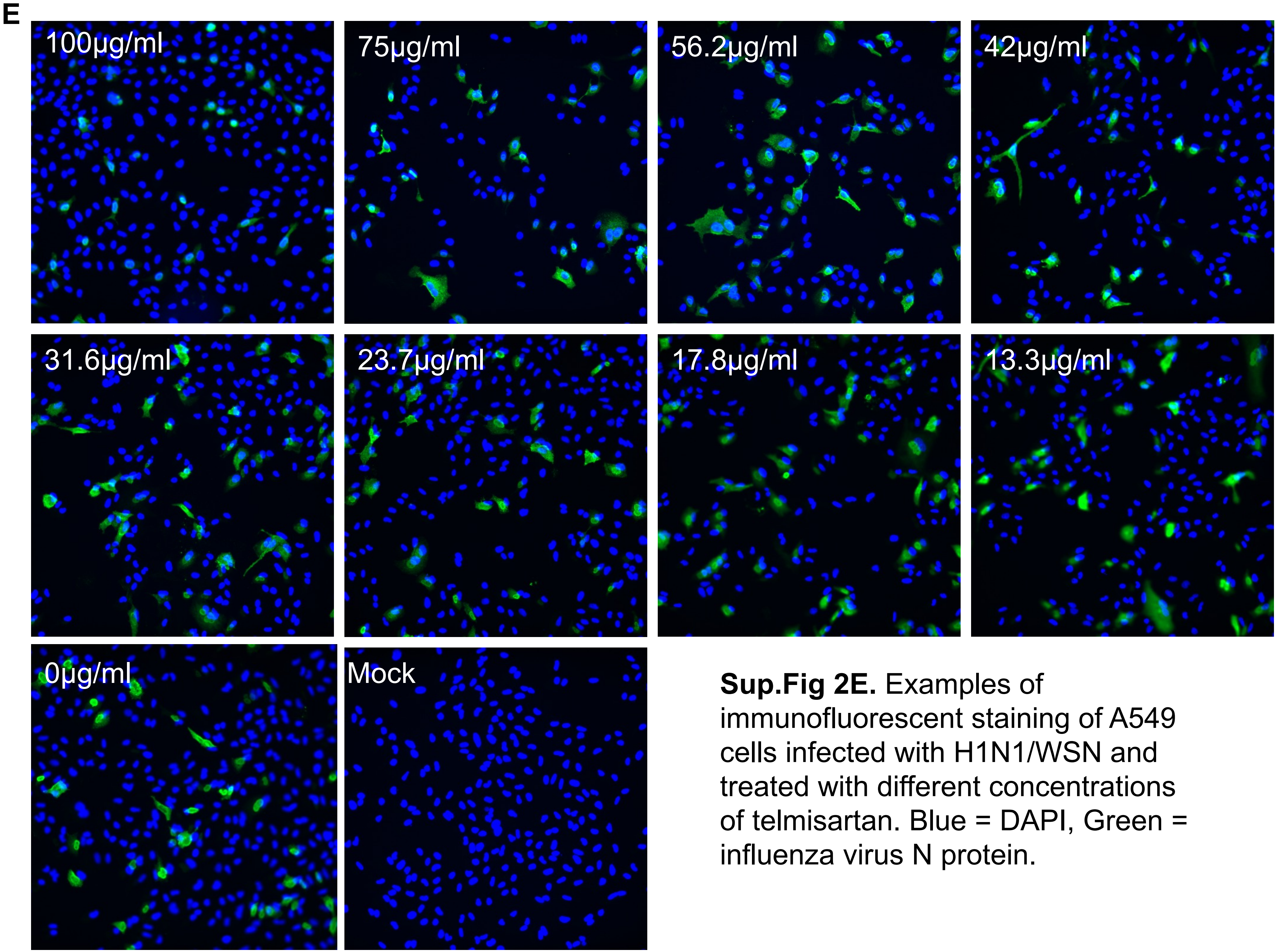


**Sup.Fig 2E.** Examples of immunofluorescent staining of A549 cells infected with H1N1/WSN and treated with different concentrations of telmisartan. Blue = DAPI, Green = influenza virus N protein.

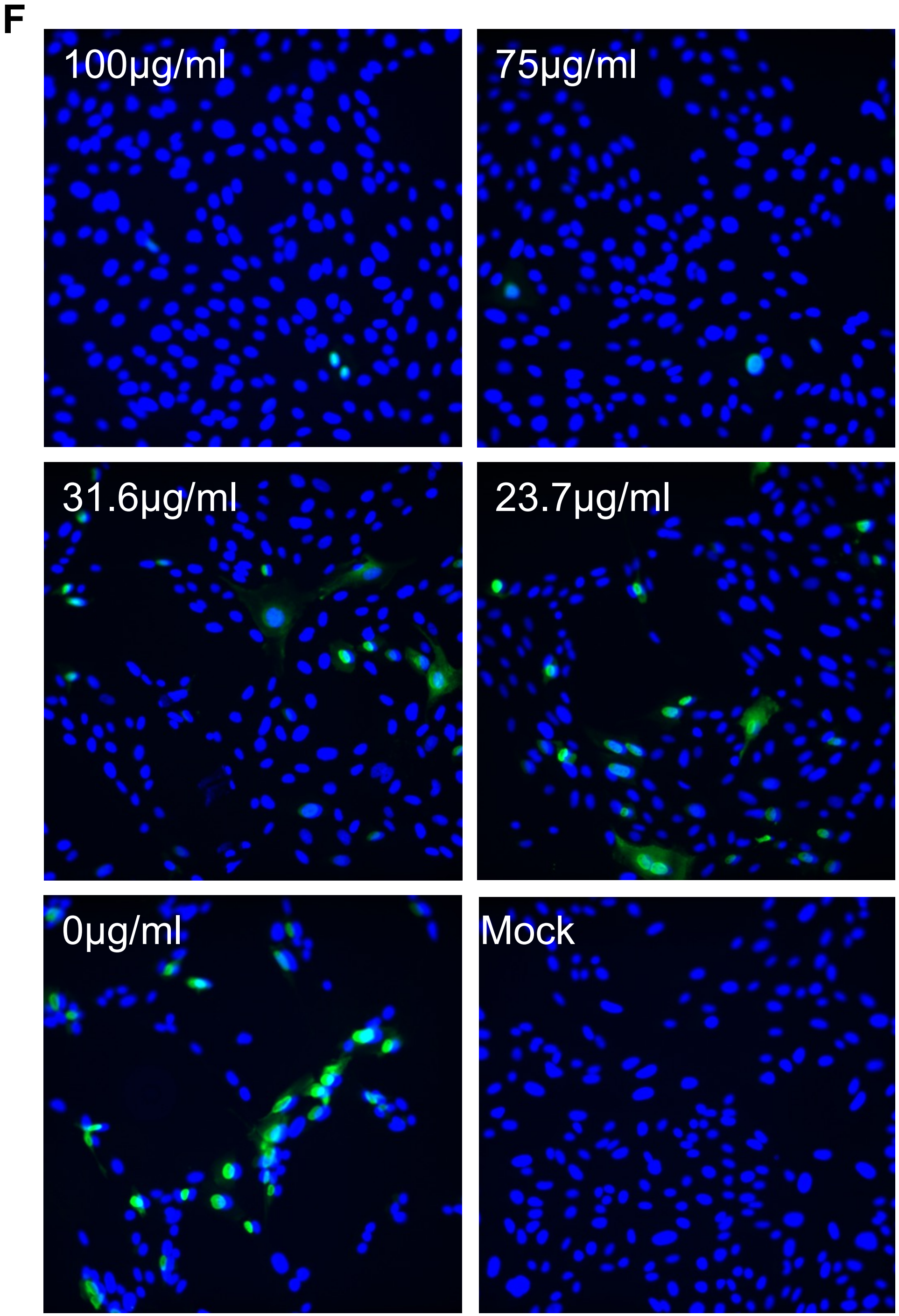


**Sup.Fig 2F.** Examples of immunofluorescent staining of MDCK cells infected with H1N1/WSN and treated with different concentrations of telmisartan. Blue = DAPI, Green = influenza virus N protein.

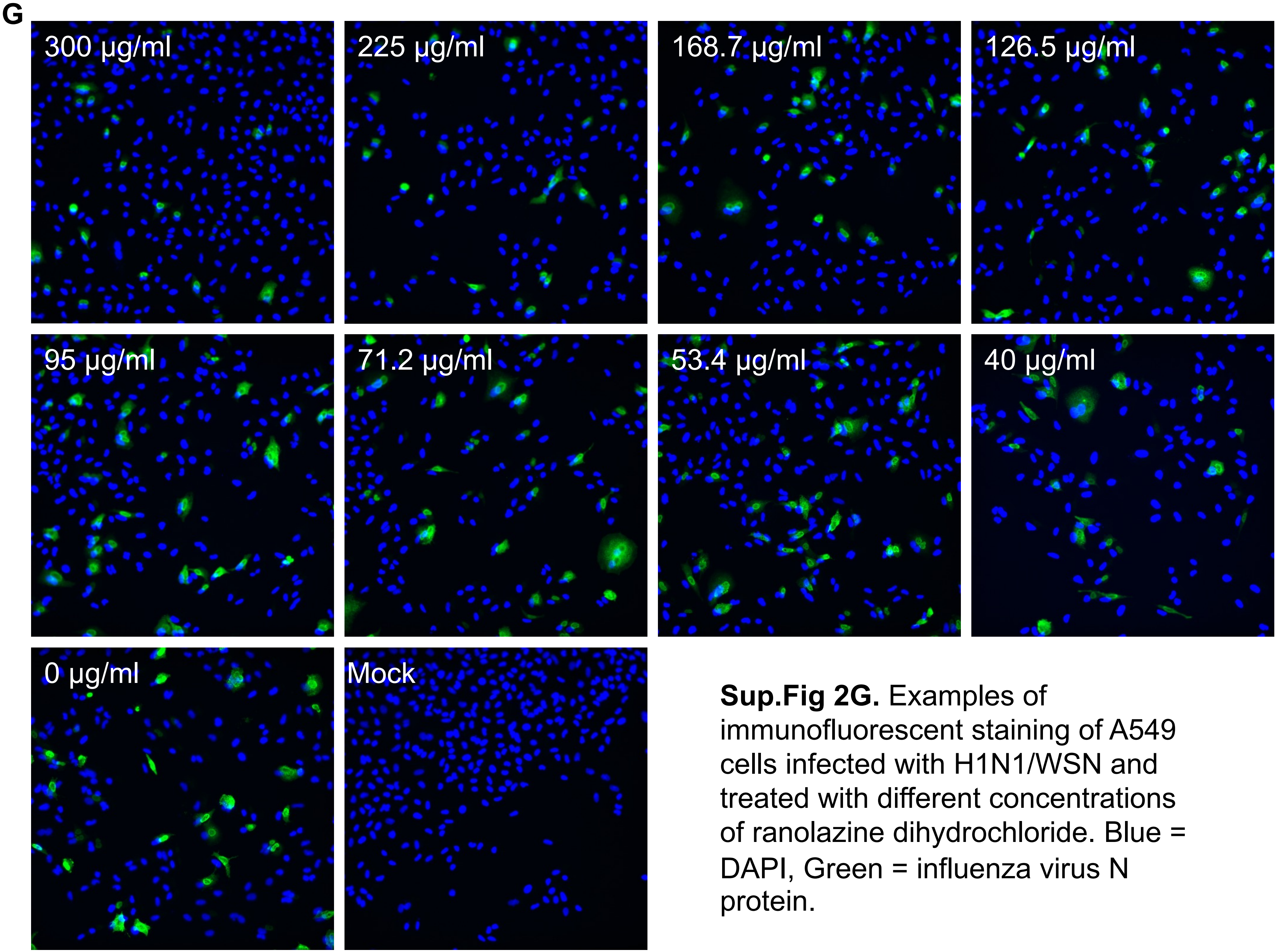


**Sup.Fig 2G.** Examples of immunofluorescent staining of A549 cells infected with H1N1/WSN and treated with different concentrations of ranolazine dihydrochloride. Blue = DAPI, Green = influenza virus N protein.

H

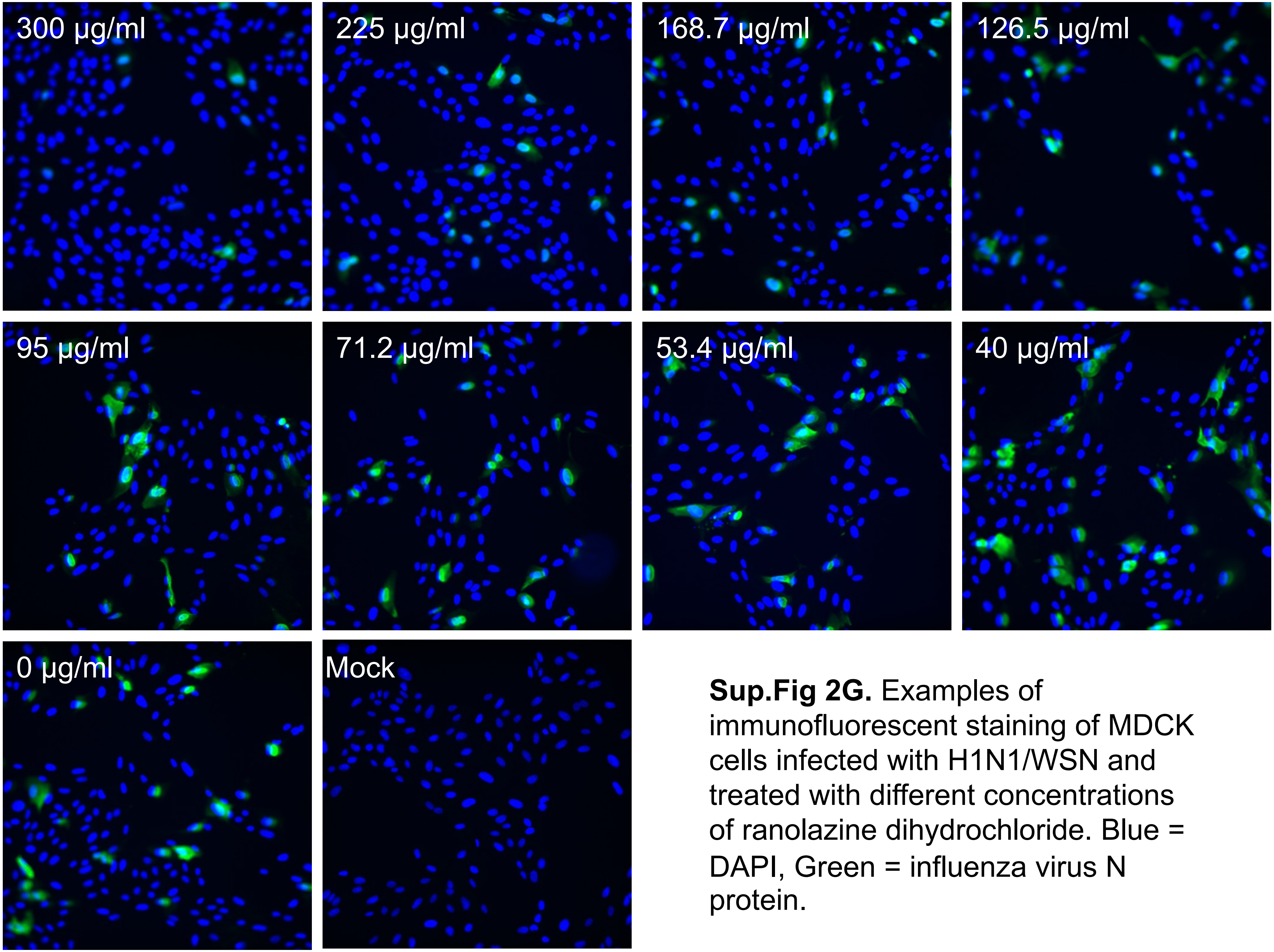


**Sup.Fig 2G.** Examples of immunofluorescent staining of MDCK cells infected with H1N1/WSN and treated with different concentrations of ranolazine dihydrochloride. Blue = DAPI, Green = influenza virus N protein.

I

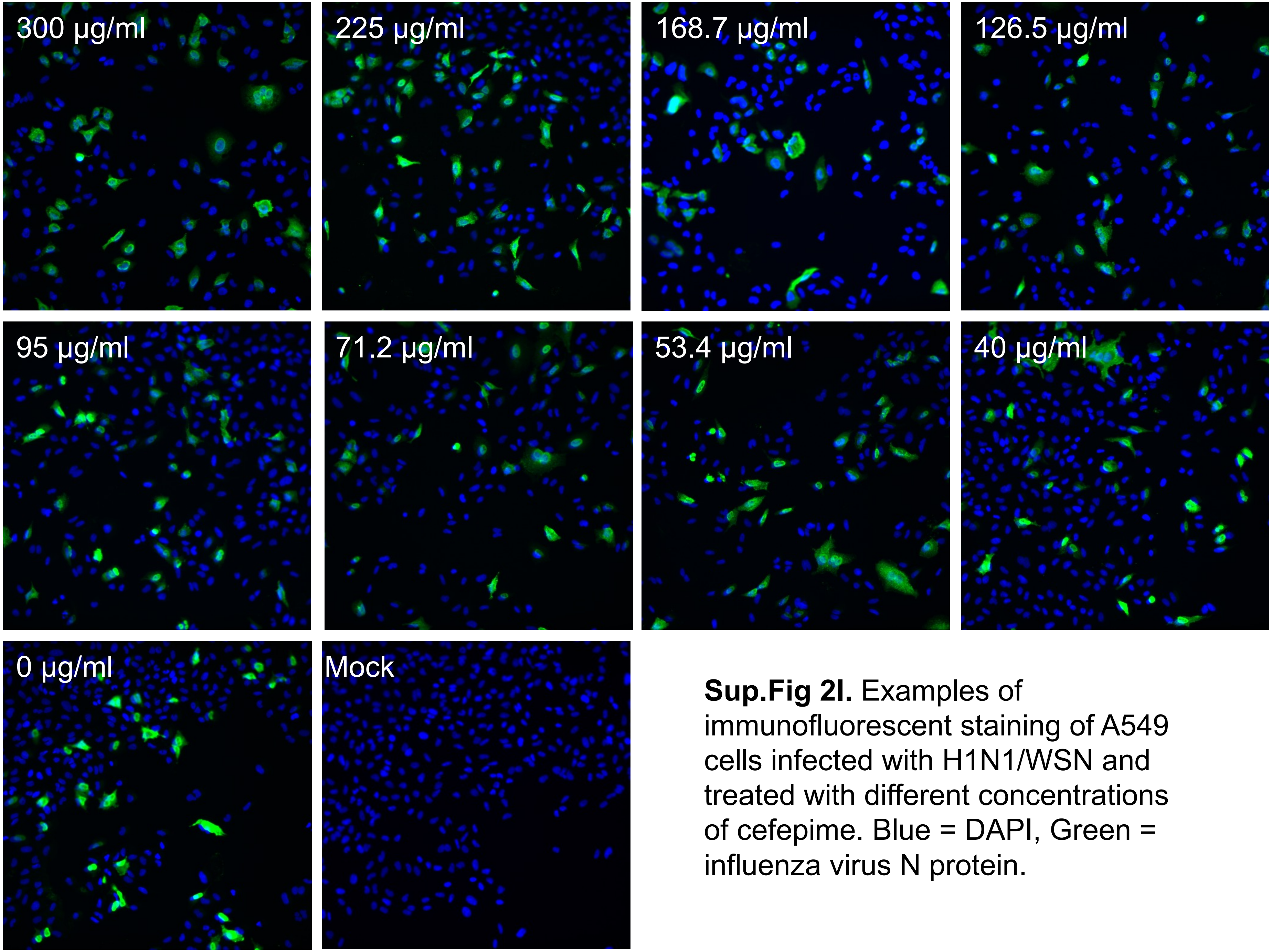


**Sup.Fig 2I.** Examples of immunofluorescent staining of A549 cells infected with H1N1/WSN and treated with different concentrations of cefepime. Blue = DAPI, Green = influenza virus N protein.

J

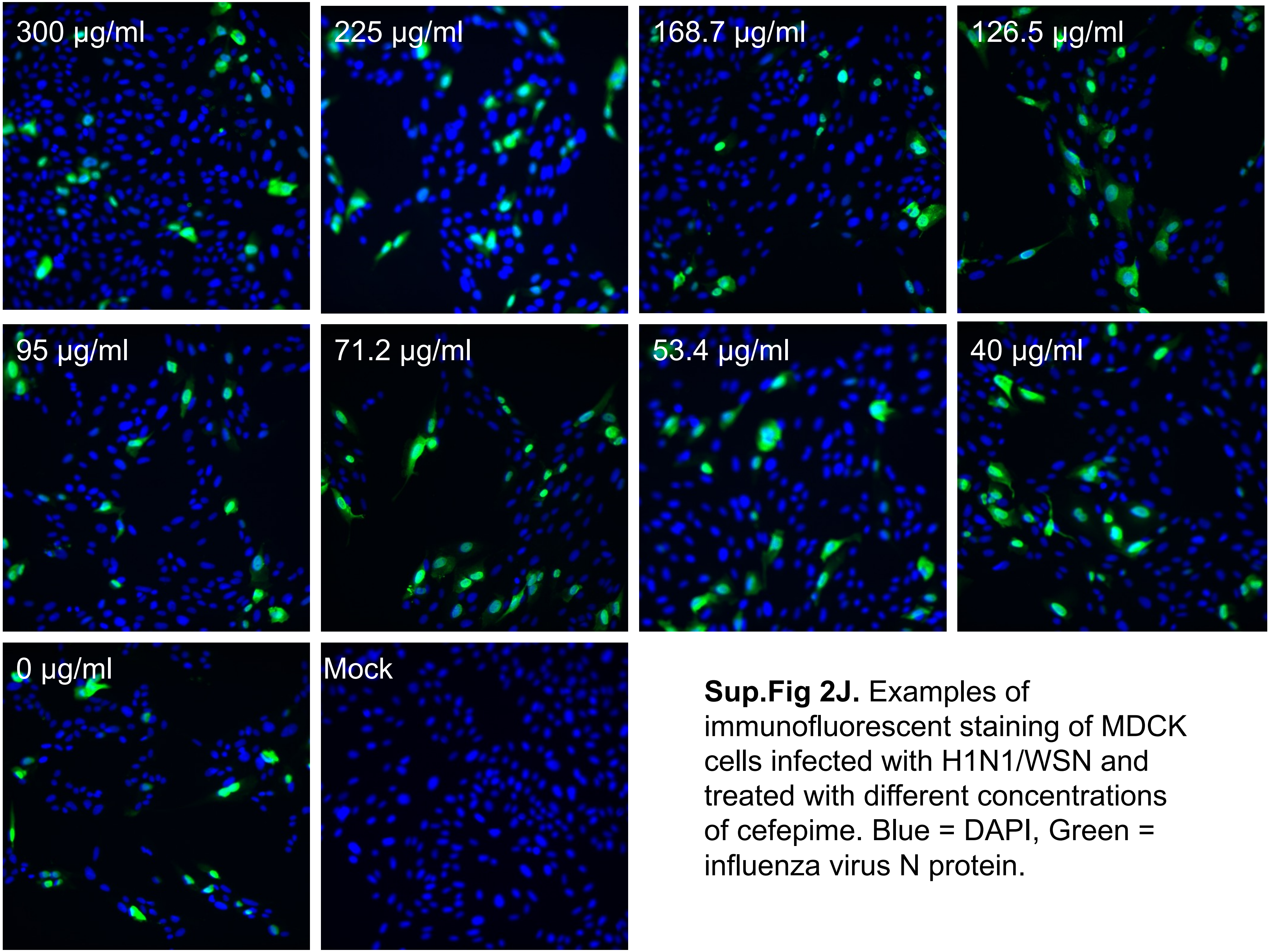


**Sup.Fig 2J.** Examples of immunofluorescent staining of MDCK cells infected with H1N1/WSN and treated with different concentrations of cefepime. Blue = DAPI, Green = influenza virus N protein.

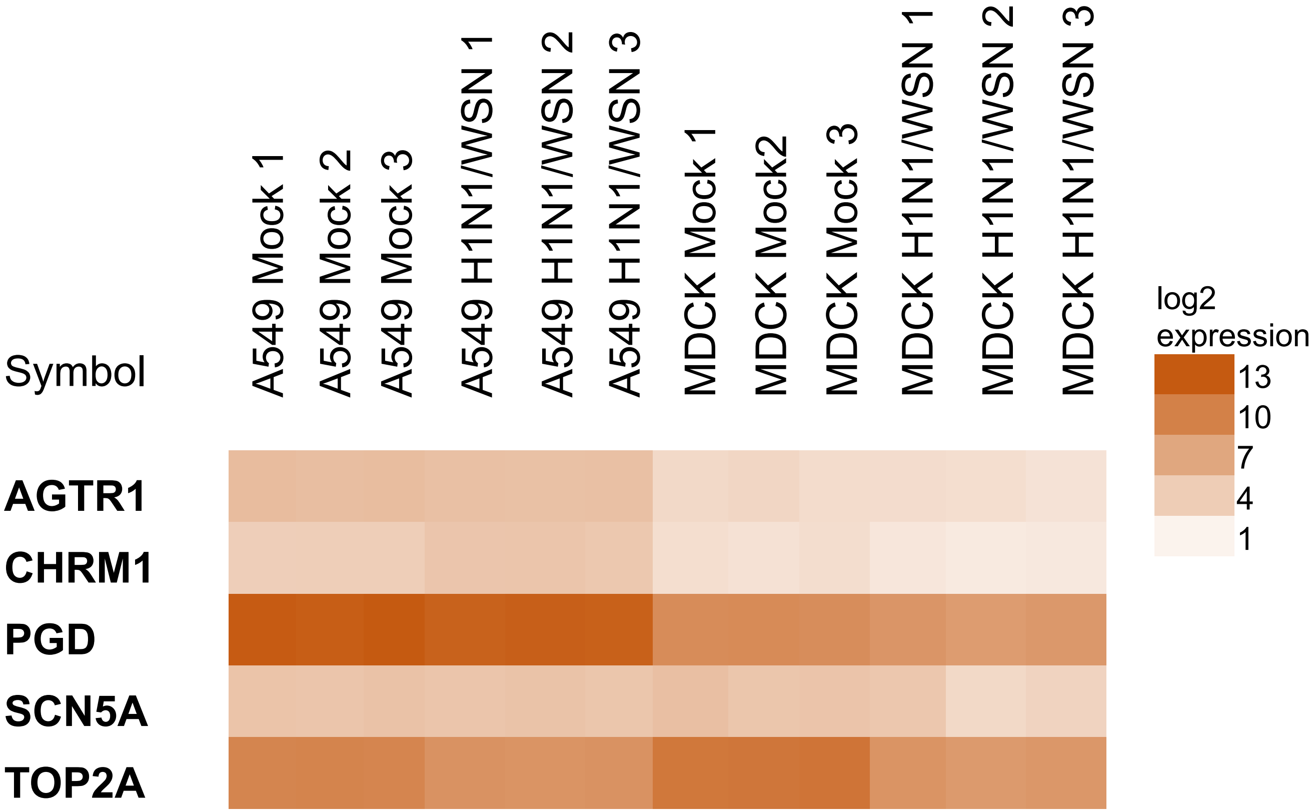


**Sup.Fig 3**. Heatmap indicating the expression of the cellular target genes in A549 and MDCK cells with and without A/WSN/1933(H1N1) infection at 10 hours post-infection. The expression data were obtained from the GEO dataset accession number (GSE31524) and normalized and $log^2$-transformed expression values were used to create the heatmap.
